# Deciphering DNA’s sequence-dependent structure and deformability with normalizing flows

**DOI:** 10.1101/2025.02.01.636041

**Authors:** Korbinian Liebl

## Abstract

The sequence-dependent structure and deformability of double-stranded DNA plays a key role in many cellular processes. Accurate description thereof has thus been a long-standing problem. Previous approaches to this problem assume a specific functional form for the elastic energy in terms of internal coordinates of the DNA double-helix. The conformational flexibility of DNA, however, is strongly impacted by several stereo-chemical effects that complicate the formulation of an accurate functional form. In this work, I propose an entirely new, AI-based method to decipher the sequence-dependent structure and deformability of double-stranded DNA. This method employs normalizing flows that capture multimodal and correlation effects between internal coordinates of the DNA double helix excellently, and hence allows one to accurately quantify deformation energies for any double-stranded DNA structure and sequence. Thus, it offers a wide range of future applications, and speaks in favor of AI-based elasticity-descriptions also for other molecules.

## Introduction

DNA encodes diverse information. Its sequence stores the chemical building plan for proteins^1 3^, but it also encodes its own deformability, which is crucial information for cellular life^4 8^. A vast number of proteins, for instance, bind to specific sequences of the DNA. This specificity often originates from slight structural and pronounced elastic differences between the DNA sequences^4 6,9^. That proteins sense DNA’s local structure and deformability and bind upon an energetically favorable response, is understood as an indirect readout-mechanism that is employed in many critical biological processes, e.g., transcription-initiation by the TATA-protein^10,11^, licensing of the origin of replication (ORC)^12^ or binding of the Tryptophan repressor protein for gene regulation.^13^

An accurate description of DNA’s structure and deformability is therefore of high relevance and has been an active field of research for several decades. A pivotal approach in this fifield has been to describe the elastic energy of DNA by a quadratic ansatz, in which the stiffness matrix can be inferred by inversion of the covariance matrix obtained from crystal structures^14^ or Molecular Dynamics simulations (MD)^15 18^. Hereby the elastic energy is conventionally expressed in dependence of rigid body transformation parameters, and recently also phosphate related degrees of freedom have been integrated.^19^

While MD simulations have greatly augmented the deciphering of DNA’s structure and deformability, they have also brought to light clear shortcomings of the quadratic ansatz: Several degrees of freedom exhibit multimodal behavior^20 24^ and for larger deformations DNA’s deformability greatly softens^25,26^. The multimodal behavior is caused by the existence of at least two dominant substates (BI and BII) in DNA’s phosphate backbone which is also evidenced by crystal structures and NMR measurements^27 30^. The difficultyin describing DNA’s deformability is further exacerbated by pronounced correlations between multiple degrees of freedom^20,31,32^. Reconciling multimodal behavior with an adapted form of the quadratic ansatz, for instance by writing the partition sum as a sum of Gaussian functions, is therefore not straightforward, as also juxtaposed DNA base-pair steps are highly cor-related and probability distributions over local segments hence not additive. Intriguingly, multimodal and correlated behavior can be unified in a multivariate Ising-model for DNA’s deformability.^31^ Such an Ising-model can be parameterized purely based on MD-data and has been shown to yield more realistic deformation energies for DNA-duplex structures and achieved excellent correlation between experimental binding affnities and computed deformation energies for the papillomavirus E2 protein complex, in contrast to the quadratic ansatz. However, the multivariate Ising-model incorporates only two substates so far and does not include the softening-behavior for larger deformations.

Although the Ising-model is still extensible in many ways, I present a fundamentally different approach to DNA’s structure and deformability in this paper. Factorizing the probability distribution for a DNA-duplex, the probability for a given structure can be computed as a product of normalizing fiow models for the local DNA segments. Normalizing fiow models are generative AI-models that invertibly map a normal-distributed latent-space to the target-space^33 35^. In this way, they offer an expression for the probability distribution of the target-space without a priori assuming a functional form. For these reasons, NF-models have been employed for various molecular-sampling problems^36 39^. In this paper, I show that an NF-based approach successfully captures the multimodal and highly correlated, sequence-dependent elastic behavior of double-stranded DNA. By parameterizing NF-models for all local sequence contexts, I have developed a general model that allows one to reliably predict deformation energies for larger double-stranded DNA structures of any sequence. The application of this model to DNA bound in a nucleosome reveals a periodic deformation pattern that arguably has strong implications on nucleosome positioning and protection of sequences that are vulnerable to UV-light.

While a few AI-models for DNA-structure prediction have already been developed^40,41^, this work presents to the best of my knowledge a first rigorous AI-model to capture doublestranded DNA’s conformational behavior accurately. It thus represents a very new concept to systematically study the mechanical genome and facilitates a plenty of future applications, as it can be used to compute deformation energies arising upon binding by proteins and in this way to quantify sequence-dependent effects. Moreover, this approach can also be extended and applied to DNA damages or modifications (e.g., cytosine-methylation) to elucidate the binding-specificity of repair proteins or energetic cost for DNA condensation, for instance. Finally, the presented approach can also be adopted for other types of molecules.

## Theory

### Design of the AI-model

I use a commonly employed geometric description for the double-stranded DNA structure. In this description, a reference frame **r**_**n**_ is assigned to each base-pair^42 44^. The parameters then follow from rigid-body transformation between successive base-pairs: **r**_**n**+**1**_ = *R*(**r**_**n**_, *dα*_*n*_, *dβ*_*n*_, *dγ*_*n*_) + *L*(*dx*_*n*_, *dy*_*n*_, *dz*_*n*_), (Fig. 1 A). Thus, each base-pair step is described by three rotational (tilt, roll, twist) and by three translational (shift, slide, rise) parameters. For the sake of simplicity, the rigid-body parameters are referred to as ‘internal coordinates’ in the rest of the paper.

**Figure 1.**
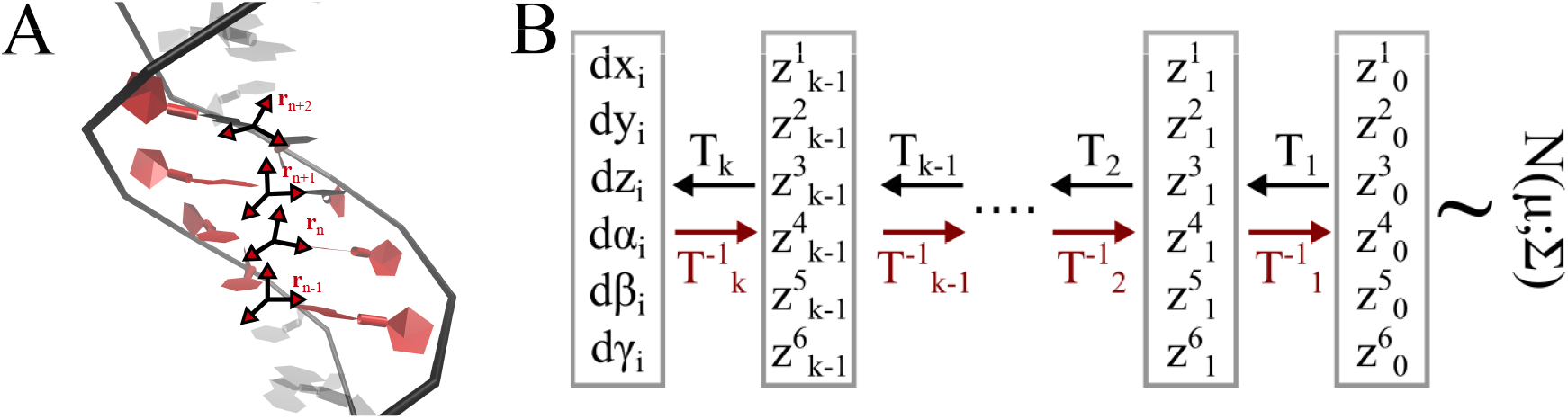
Parameterization and description of the dsDNA structure and conformational fiexibility. (A) The structure of a dsDNA molecule is parameterized by rigid-body transformations, where a axis-system is spanned on each base-pair step (**r**_**i**_). The parameters describe rotational and translational functions that map the axis-system of a base-pair step to the adjacent axis-system. (B) Normalizing flow models map from a normal-distributed latent-space to the parameters that describe the dsDNA structure. Iapping is achieved through multiple layers and is invertible.

As shown in Fig. 2, and noted in earlier works on DNA deformability, the fluctuations of these internal coordinates can be highly correlated. The correlations, however, decay significantly over distance, so that beyond nearest-neighbor effects can be neglected, i.e., internal coordinates of a base-pair step are correlated only to themselves and their juxtaposed base-pair step^20,31,32^. The probability for a given N base-pair-steps long double-stranded DNA structure can hence be written as:

**Figure 2.**
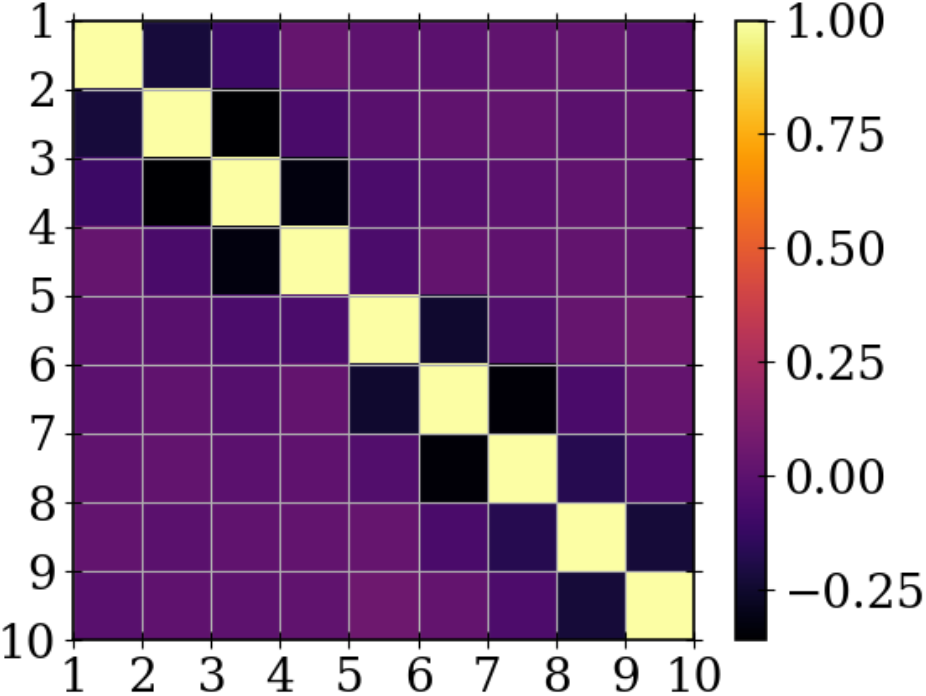
Color-coded correlation matrix for the twist-parameters of successive base-pair steps obtained from MD simulations. The near-diagonal elements show noticeable anti-correlation (black squares). Remaining elements indicate negligible correlation (purple squares). Thus, an accurate model for dsDNA’s elasticity must capture nearest-neighbor correlations, whereas long-ranged correlations may be neglected.

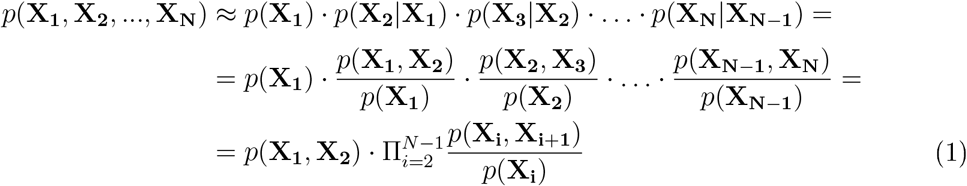

Now, the probability distributions of the local segments can be trained with Normalizing Flow models, *p*(**X**_**i**_) ≈ *p*^*NF*^ (**X**_**i**_) and *p*(**X**_**i**_, **X**_**i**+**1**_) ≈ *p*^*NF*^ (**X**_**i**_, **X**_**i**+**1**_). Normalizing Flow models are generative AI-models with an invertible transformation between latent-space (**z**) and internal coordinate-space (**X**), Fig 1B. In this way, the likelihood of the data can be computed,

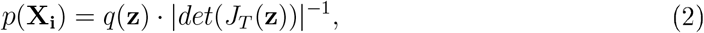

where T denotes the transformation from latent to coordinate-space. This transformation is typically trained as a sequence of transformations (*T* = *T*_*k*_ ∘ *T*_*k*−1_ ∘ … ∘ *T*_1_), so that the log-likelihood becomes:

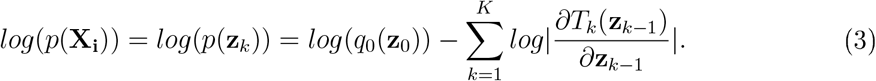

Note that *q*_0_(**z**_0_) is the base-distribution, which is described as a multivariate normal distribution in the following. The parameters of the Normalizing fiow models are trained by minimizing the forward Kullback-Leibler divergence *L*(*ϕ, θ*) = *D*_*KL*_ [*p*^*^(**X**_*i*_)||*p*(**X**_*i*_; *ϕ, θ*)]^33,35^. *ϕ* are the trainable parameters of the base-distribution, and *θ* of the transformation from latent-to coordinate-space (*T* (*θ*)). The gradients of the loss-function with respect to the trainable parameters can be estimated as:^35^

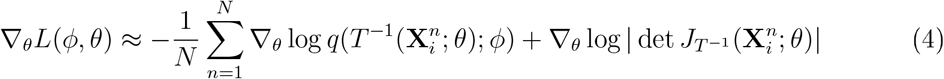

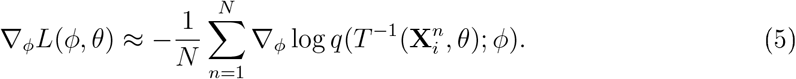

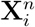 are the samples for base-pair step i obtained from atomistic MD simulations prior to training the NF-models. The transformation in the NF-models consists of normalization layers (assuming a fixed statistics) and real NVP transformations^35,45^. In the latter, data of dimension D is split into two channels (1:D/2, D/2+1:D), and transformed as follows:

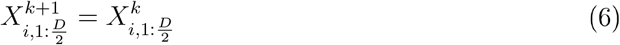

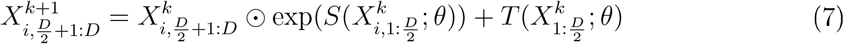

This design triangulates the Jacobian-matrix and the scaling- and translation-functions 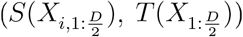 are in this work described by three layers of tanh-operations. More details on the architecture of the NF-models are given in the Supplement. The NF-models were developed with pytorch, and training was performed for 20000 epochs with the adam-optimizer.^46^ The learning-rate was decreased from 0.02 to 0.002 during training.

### MD simulations

A 16bp-long dsDNA structure of complementary sequence (5’-GCGCAATGGAGTACGC-3’) was generated with the nab-module of amber and used as starting structure for the MD simulation.^47^ The Tumuc1 force field was used to describe the DNA-molecule^48^ and the TIP3P water-model with a salt concentration of 250mM NaCl was employed.^49^ The system was energy-minimized with the steepest descents optimizer in 5000 steps. Subsequent equilibration was performed for 250,000 steps with a time-step of 2fs in the NVT-ensemble using the Berendsen thermostat with a reference temperature of 303.15 K and a coupling constant of 1ps.^50^ In the equilibration, positional restrains with a force-constant of 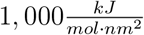 for each dimension. The equilibrated structure was used as input to the data-gathering simulation of 25,000,000 steps with a time-step of 2fs (without any restraints). The simulation time for the data-gathering simulation is therefore 50ns. As shown in ref^31^, this is sufficient to converge the fluctuations of the internal coordinates of a dsDNA molecule. Coordinates were written out every 5,000 steps.

Three further complementary dsDNA sequences were simulated (5’-GCAAACAAGCCCGACCCG’-3’, 5’-GCAATAACGCACTAGAACG-3’, 5’-GCAGGATGGAGTACTGACG-3’) in the same way as the previous sequence, except that data-gathering simulations lasted 100ns. Note that these three sequences cover all possible double base-pair steps (10) as well as all possible triple base-pair steps (32). All simulations were carried out with the Gromacs package.^51^ The geometries of the sampled DNA structures were parameterized with the Curves+ software^44,52^.

## Results and Discussion

### Comparison between MD- and NF-ensemble

The log-likelihood for a given local conformation can be evaluated with Normalizing Flows according to equation 3. Besides, new conformations can also be generated by drawing samples from the base-distribution and transforming them to the internal coordinate-space. In the following, I compare the distributions generated with Normalizing Flows to those obtained from MD-simulations, and in this way discuss the accuracy of the model. The distributions of the internal coordinates for various sequences are shown in Fig 3, 4 and Fig S1-S20 (grey: MD-sampled, green: NF-generated). The distributions of several internal coordinates show distinct anharmonic behavior, which is excellently reproduced by the NF-models. In fact, one might also intuitively expect anharmonic behavior of the coordinates. At short-distances in direction of the helical axis (quantified by the rise-parameter), for instance, the base-pairs experience strong repulsion, but the restoring force for larger separations should vanish. Thus, a Morse-oscillator is a more plausible assumption for the elastic energy than the harmonic approximation, which represents only a first-order approximation to it. However, the elastic behavior of the DNA double-helix is further complicated by backbone related substates and substantial correlation between the internal coordinates. As shown in Fig 3 and 4, deviations from Gaussian distributions (and hence from a harmonic energy landscape) are especially visible in the shift, slide and twist parameters, and the NF-models are able to capture multimodal behavior. Recall from equation 1 that two types of NF-models need to be trained, for single *p*(**X**_*i*_) and double base-pair steps *p*(**X**_*i*_, **X**_*i*+1_), and both types of models display excellent description of the conformational fluctuations.

**Figure 3.**
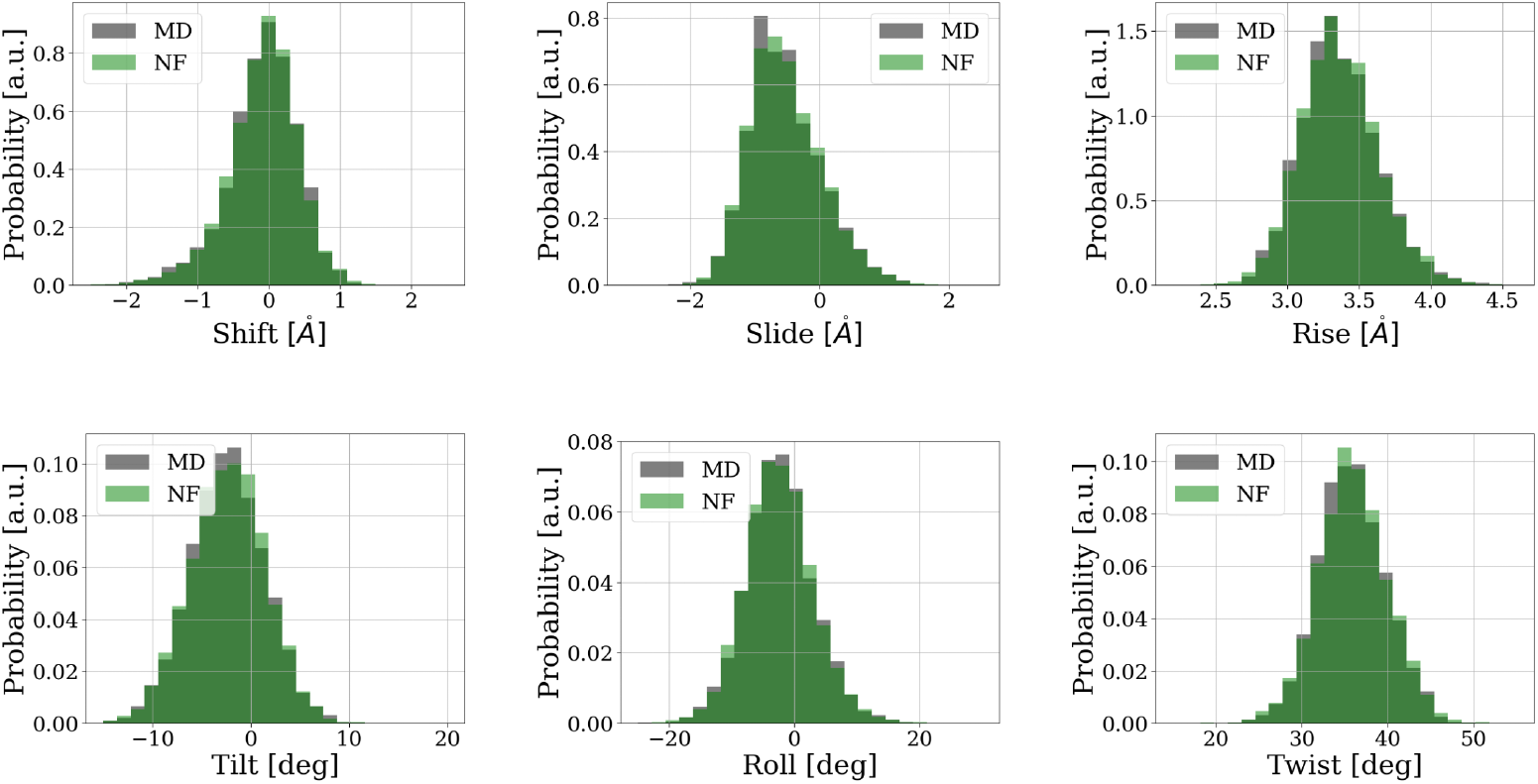
Distributions of the internal coordinates for the 5th base-pair step as sampled from atomistic MD simulations (grey) and the Normalizing Flow double-step model (green).

**Figure 4.**
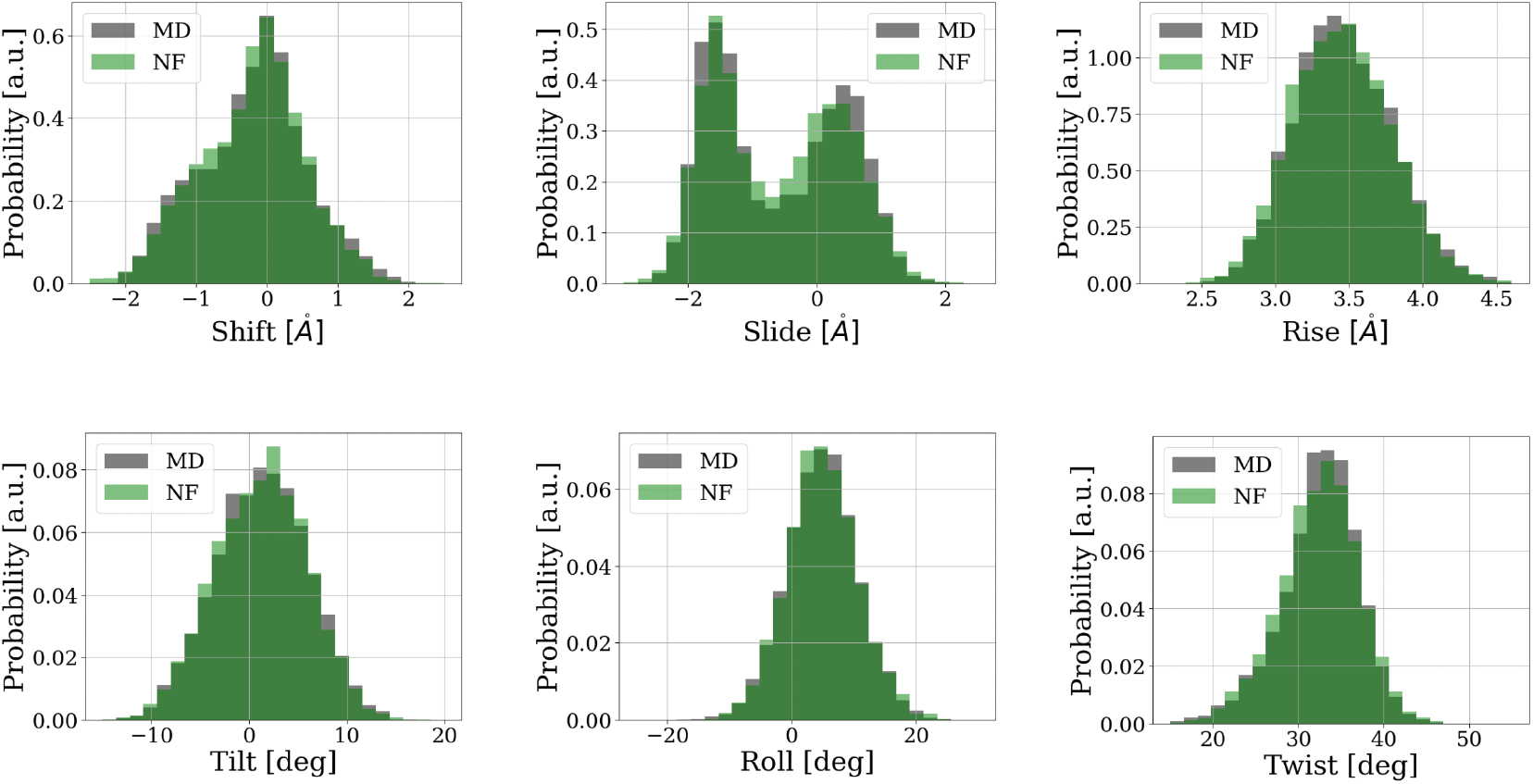
Distributions of the internal coordinates for the 8th base-pair step as sampled from atomistic MD simulations (grey) and the Normalizing Flow double-step model (green).

Moreover, the NF-models not only capture the 1d-distributions accurately, but also reproduce correlations between the internal coordinates excellently. Fig 5A depicts the correlations between the internal coordinates of one base-pair step sampled from atomistic MD simulations and reveals an intricate conformational behavior, as the correlation between internal coordinates varies greatly. The slide and twist parameters, for instance, are strongly positively, but slide and rise are negatively correlated. In previous attempts to model the deformability, these effects have been directly incorporated by computing a stiffness-matrix as inverse of the covariance matrix of the internal coordinates. As shown in Fig 5B, however, NF-models also capture the correlations in quantitative agreement with the atomistic dataset. By accurately reproducing the structural distributions and correlations, NF-models yifield an overall excellent agreement with MD-data and accomplish what has been a major challenge for the description of dsDNA’s deformability so far.

**Figure 5.**
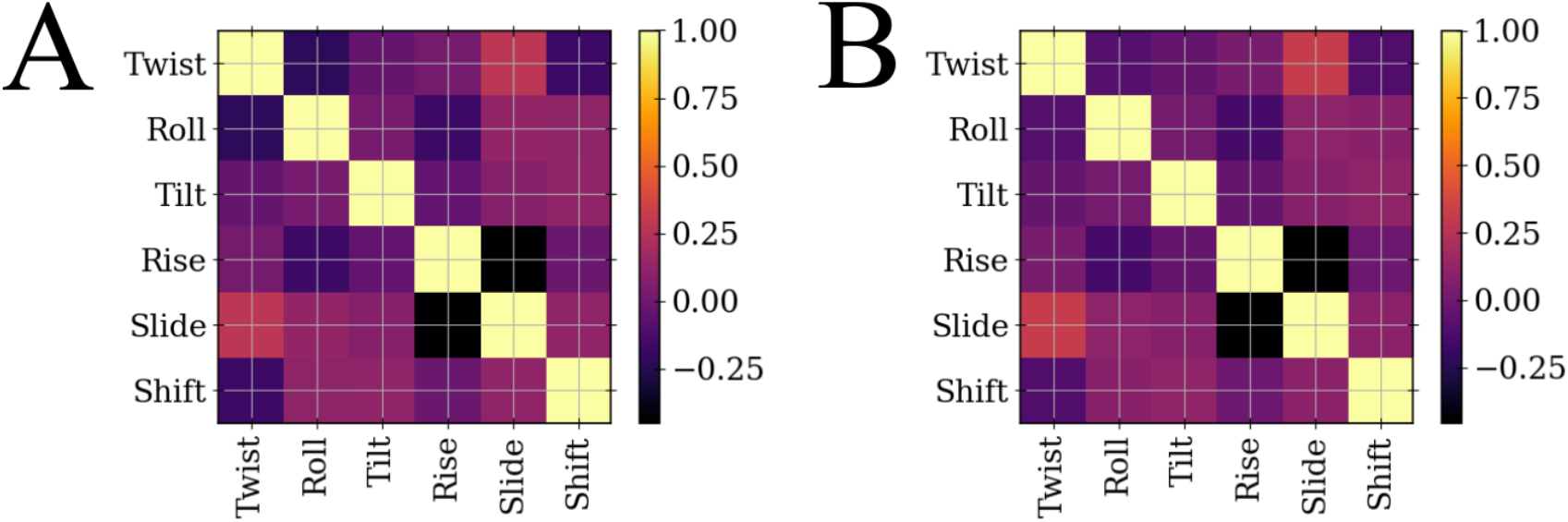
Correlation matrices between the six parameters of one base-pair step, sampled by MD-simulations and NF-models (A,B).

### Evaluation of deformation energies

Having trained the NF-models for the various base-pair steps and double steps, the deformation energy can be evaluated for any given conformation (equation 1). In this way, I have computed deformation energies for the entire trajectory of the test-sequence. Only the central ten base-pairs were considered in the energy-evaluation to filter out the impact of terminal fraying events, resulting in 54 internal coordinates. Additionally, I have also computed the deformation energy for the mean-structure (consisting of the time-averages of all parameters). The di fferences between the frame-wise energies and the energy for the mean-structure (*E*_*mean*_ = 4.2*kcal/mol*) are plotted in Fig 6. Note that only a few of the sampled structures have a lower energy than the mean-structure. In other words, the NF-based model recognizes the temporally averaged structure as energetically favorable, but it does not have to be the native state due to multimodality. This further indicates a realistic description by the NF-based model which is also confirmed by classifying the low- and high-energy samples. All of the samples that are evaluated as energetically unfavourable show clear structural deformation, such as local bending and kinking (snapshots 2,7 in Fig 6) or undertwisting (snapshots 3,5). On the other hand, structures that are predicted to be energetically favorable (snapshots 1,4,6), appear as regular B-DNA duplex structures. Intriguingly, the model predicts that distortions in the DNA-duplex structures are equilibrated rapidly, as a strongly deformed structure (snapshot 5) can relax within a few picoseconds to an intact shape again (snapshot 6). Further studies of such relaxation processes or hypothetical deformation solitons along dsDNA molecules are of great interest, and can be addressed reliably with the proposed NF-based model.

**Figure 6.**
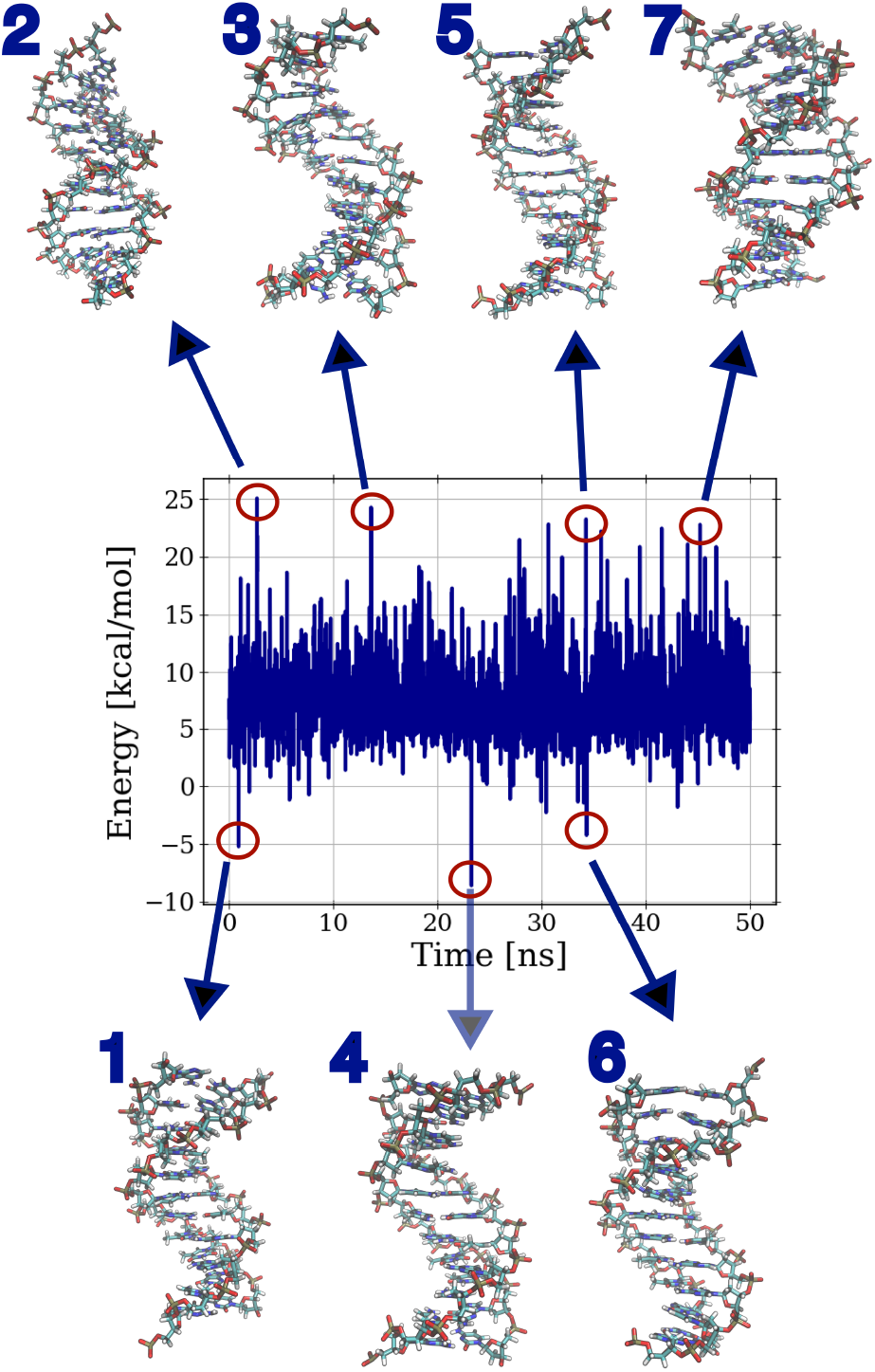
Deformation energies calculated for the different structures sampled during simulation. Note that the plotted values include subtraction of the energy for the mean-structure. Snapshots 1,4,6 depict energetically favorable states, and snapshots 2,3,5,7 strongly deformed structures.

### Generalization of the model

The analysis and application so far concerned only one dsDNA sequence. To develop a general model that captures the entire sequence space, three additional DNA duplexes with different sequences have been simulated. These three sequences capture the entire sequence space of double or triple base-pairs. Thus, by training the NF-models for the three simulations, the structure and deformability of the entire ‘DNA-alphabet’ can be modeled. Comparison of some of the resulting models to MD-data is given for a few selected sequences in Fig S21-S24. This generalizing approach allows one to compute deformation energies for any given DNA-duplex structure, and therefore enables many future applications. For example, one can compute deformation energies also for DNA molecules bound by proteins. In this way, sequence-dependent energetic costs upon protein binding (indirect readout) can be systematically calculated, hence enabling a better, quantitative understanding for sequence-specific recognition mechanisms utilized by proteins. Moreover, one may not always be interested in the deformation energy computed over one longer DNA-segment, but rather calculate individual deformation energies for local sites to generate a stress-profile along the DNA-molecule. Fig 7 shows such an application for dsDNA in the nucleosome complex. The deformation-profile along the sequence (Fig 7B) does not show significant outliers, but most of the values are in an reasonable range (0-10 kcal/mol), which further confirms that the NF-models for the sequences have been trained appropriately. In fact, the energy-profile follows an intriguing, oscillating pattern that is more clearly revealed in the energy-autocorrelation function (Fig 7C). The autocorrelation-function shows peaks in multiples of the number of base-pairs per DNA-turn (∼ 10.5), and minima for half-integer multiples. Such a pattern has already been observed in earlier studies, although mostly employing simpler models with less degrees of freedom^53,54^. Overall, the NF-based deformation energy profile is hence quantitatively and qualitatively reasonable. Such insight into nucleosomal deformations is of high relevance, as it greatly impacts nucleosome positioning along the genome^55,56^. Also, local, conformational fluctuations of nucleosomal DNA play an important role in many genetic processes, as they signal access to a larger number of histone-modification proteins, for instance^57,58^. In addition, it is well understood that the UV-damage pattern of nucleosomal DNA also follows an oscillatory pattern, and that UV-vulnerable sequences (predominantly TpT-steps) are placed at sites where their deformation dispels an excitation required for UV-damage formation^59 62^. Thus, it is tempting to speculate that DNA’s sequence-dependent stiffness encodes an intrinsic UV-light protection-mechanism activated upon nucleosome formation.

**Figure 7.**
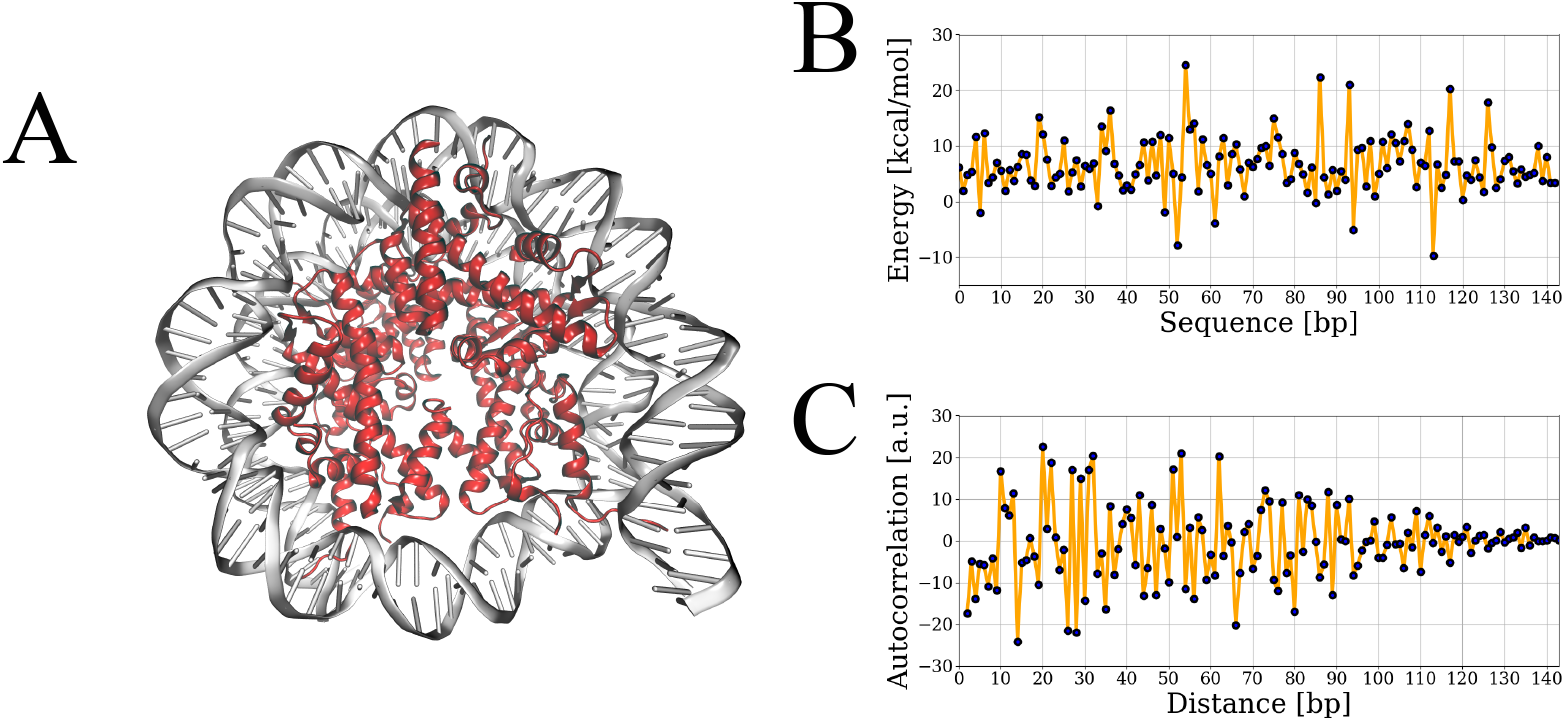
Evaluation of deformation energies for nucleosomal DNA.^63^ (A) Complex-structure, (B) Deformation energy profile along the sequence (subtracted by energy-terms for the mean-structures), and (C) Autocorrelation function for the sequence-dependent deformation energies.

## Conclusions

DNA undergoes significant sequence-dependent deformations in a vast number of cellular processes. Accurate quantification of the energetic costs is therefore paramount to understanding how these processes work. Previous attempts to model DNA’s deformability have relied on specific assumptions of the functional form. Here, a harmonic ansatz for the functional form, for which stiffnesses are derived from inversion of sampled covariances, represents the most frequently discussed and applied approach. This ansatz, however, shows substantial limitations, as the conformational behavior of dsDNA is impacted by several steric effects. For instance, the harmonic approximation overestimates energetic costs for larger deformations^25,26^ and neglects pronounced multimodal behavior induced by the phosphate backbone^20 24^.

In this paper, I have developed an entirely new, AI-based approach to decipher dsDNA’s sequence-dependent structure and deformability. The conformations of base-pair steps are described by normalizing flow models, which yifield an expression for the distributions of the internal coordinates through invertible mapping from a normal-distributed latent-space. Thus, the normalizing flows importantly do not a priori assume a specific functional form for dsDNA’s deformability. This represents a substantial advantage over previous approaches, and empowers the NF-models to capture the multimodal distributions of the internal coordinates as well as correlations between them in excellent agreement with atomistic MD simulations.

By training NF-models for all double- and triple-base-pair sequences, I have developed a consistent set of NF-models that allows one to calculate deformation energies for any given DNA duplex structure or sequence. Application on structures sampled during a MD simulation indicates robust performance of the developed NF-based model; high-energy states could have been identified as visibly deformed structures, whereas low-energy states correspond to native-like B-DNA duplex structures. In addition, the computed deformation-energy profile for the nucleosome structure shows a periodic pattern that agrees with earlier reports^53,54^ . In conclusion, the energy-calculations with the NF-based model on the two systems combined with the comparisons of NF-sampled distributions to MD distributions indicate a high level of accuracy. Still, further extensions may be considered in future studies. The presented model neglects any correlation beyond nearest-neighbors, but given the excellent reproduction of MD distributions, NF models may also work well for higher dimensions and allow one to also include next-to-nearest-neighbor correlations. Possibly, the largest limitation of the model results from inaccuracies of recent atomistic force fields. The stacking-interactions between base-pairs show substantial disagreement with experimental data,^64^ and also the puckering of the sugar-moieties is still debated^65 67^. These deficiencies in the description of molecular interactions is likely causative to reported inaccuracies for elastic-moduli of dsDNA^68,69^. It is hence also of interest to train the NF-models on crystal- and NMR-structures depositited in the pdb-database, i.e., to train based on experimental and not on MD-data.

Altogether, the NF-based approach facilitates a wide range of applications. It can be used to screen the energy-landscape for nucleosome-positioning along the entire genome, or any other protein/dsDNA complex. In this regard, NF-models may also be trained for DNA-damages or mismatches, in order to assess to what degree repair proteins detect defects by sensing the local deformability. Another important extension are methylated cytosine sequences to get a better understanding how epigenetic modifications impact structure and deformability. Finally, this normalizing fiows based approach can also be adopted for other molecules or lattices. It seems plausible that the presented framework may work generally well for polymers, and can therefore be analogously adopted for dsRNA, for instance.

## DATA AND CODE AVAILABILITY

The NF-models and relevant input-files are accessible at 10.5281/zenodo.14783005.

## ACKNOWLEDGMENTS

This project received funding from the European Union’s Framework Programme for Research and Innovation Horizon Europe (2021-2027) under the Marie Sk1odowska-Curie grant agreement no. 101109916. Computational resources were provided by the University of Chicago Research Computing Center and the NIH-funded Beagle-3 computer (NIH award 1S10OD028655).

## AUTHOR CONTRIBUTIONS

K.L. designed the research, performed simulations, developed the software, analyzed the data, prepared the figures, and wrote the manuscript.

## SUPPORTING INFORMATION

Supporting material includes information on the normalizing fiow models and comparisons between the ensembles sampled by MD and the normalizing fiow models.

