## Supplement Information for "Deciphering DNA’s sequence-dependent structure and deformability with normalizing flows"

Korbinian Liebl*

*Chicago Center for Theoretical Chemistry, Institute for Biophysical Dynamics, and James Franck Institute, Chicago*

#### Architecture of the Normalizing Flow models

The architecture of the trained Normalizing Flow models consists of six real-NVP transformations and three normalization layers. Each of the six real-NVP transformation consists of three layers, where the scaling and translation functions are element-wise tanh-operations:

$$S(\mathbf{X}) = \mathbf{B}\tanh(\mathbf{A}\mathbf{X}) \tag{1}$$

$\mathbf{A}$ and $\mathbf{B}$ are matrices composed of trainable parameters.

Normalization layers have the form:

$$N(\mathbf{X}) = \mu + (\frac{\mathbf{X}-\beta}{\alpha}) \odot \sqrt{\sigma^2+\epsilon} \tag{2}$$

Inverse normalization is then given by:

$$N^{-1}(\mathbf{X}) = \alpha \odot \frac{\mathbf{X} - \mu}{\sqrt{\sigma^2 + \epsilon}} + \beta \tag{3}$$

Note that mapping from the coordinates to latent-space follows in inverse direction. The probability distribution for the coordinates also depends on the Jacobian-determinant. In theory, $\sigma$ and $\mu$ are not constant, but depend on the training-samples. Capturing this dependence in the Jacobian would be computationally highly inefficient. It is therefore common to assume that $\sigma$ and $\mu$ are constant ('fixed statistics'). This approximation becomes more accurate with increasing sample-size.

### Comparison of MD- and NF-ensemble

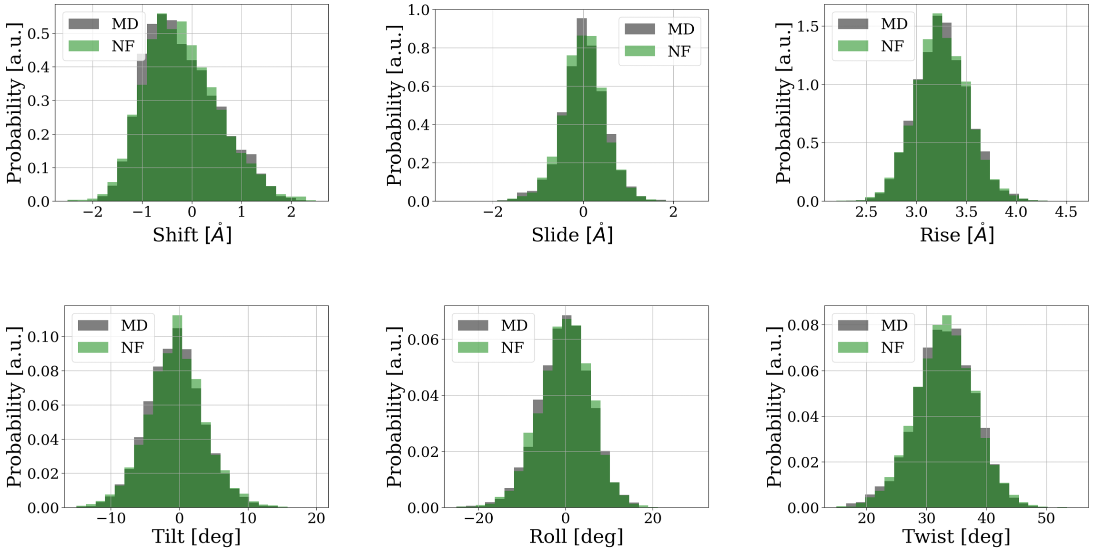


Figure 1: Distributions of the internal coordinates for the 3rd base-pair step as sampled from atomistic MD simulations (grey) and the Normalizing Flow double-step model (green).

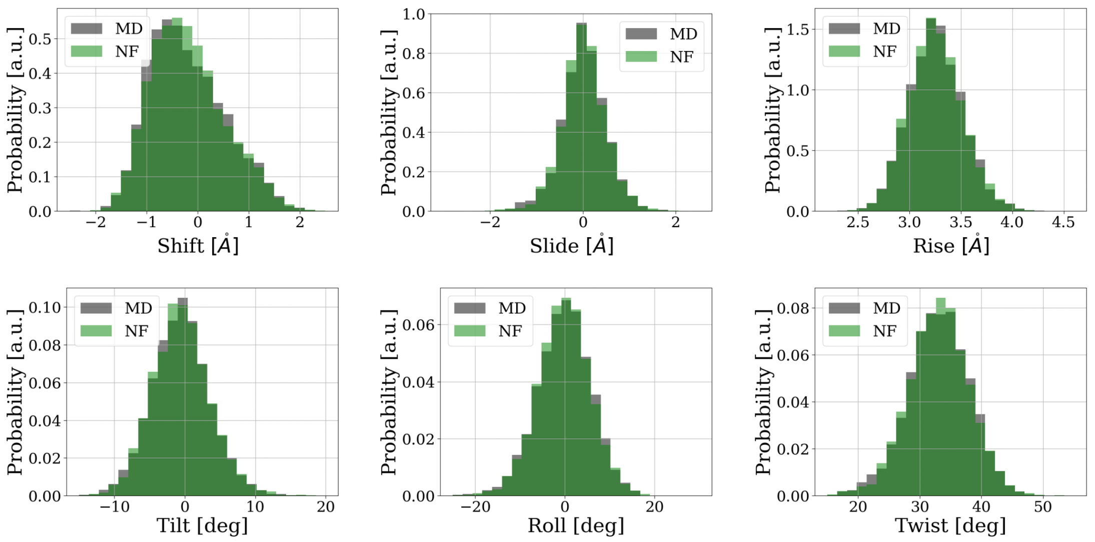


Figure 2: Distributions of the internal coordinates for the 3rd base-pair step as sampled from atomistic MD simulations (grey) and the Normalizing Flow single-step model (green).

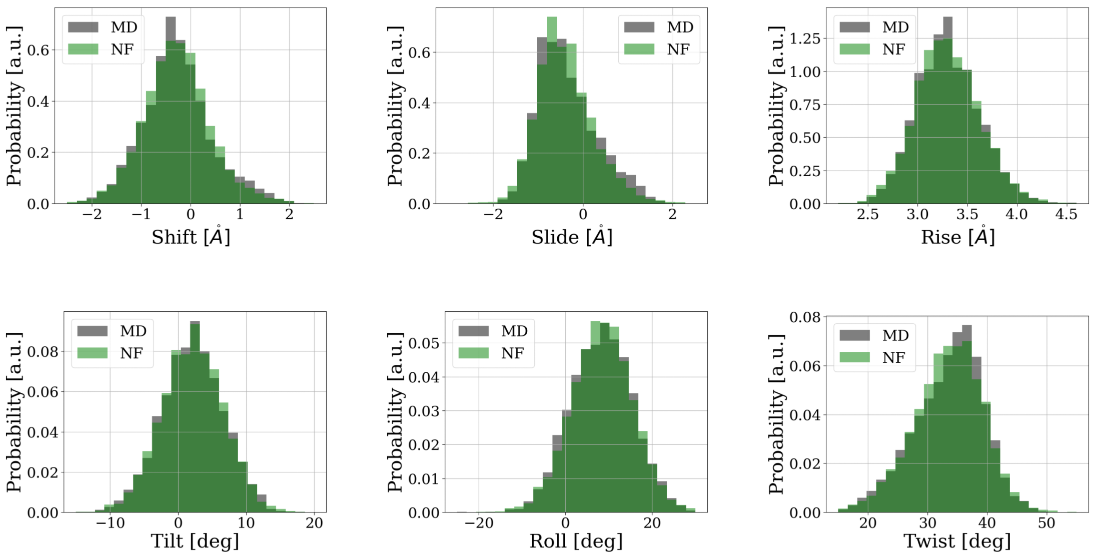


Figure 3: Distributions of the internal coordinates for the 4th base-pair step as sampled from atomistic MD simulations (grey) and the Normalizing Flow double-step model (green).

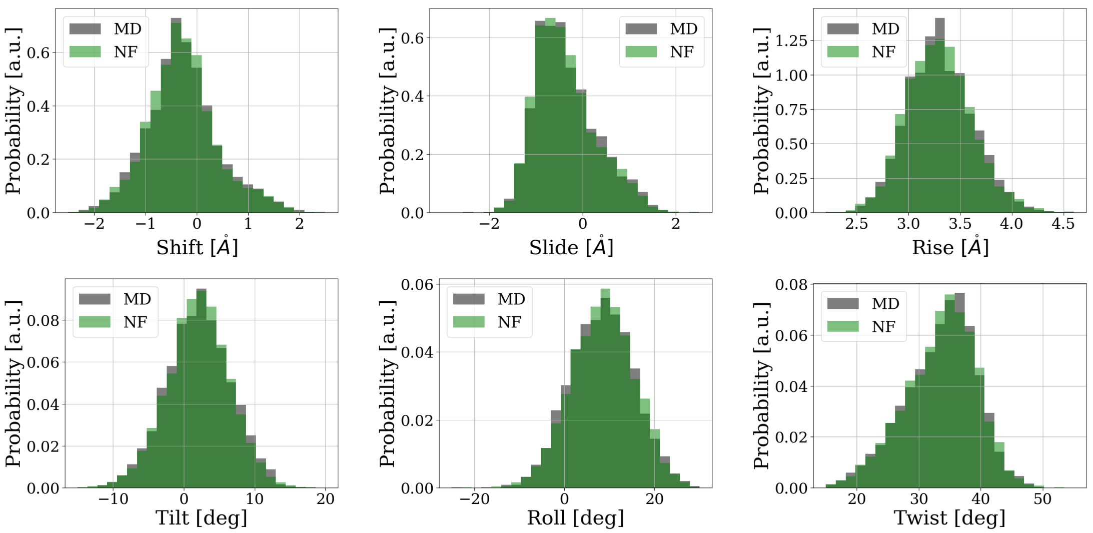


Figure 4: Distributions of the internal coordinates for the 4th base-pair step as sampled from atomistic MD simulations (grey) and the Normalizing Flow single-step model (green).

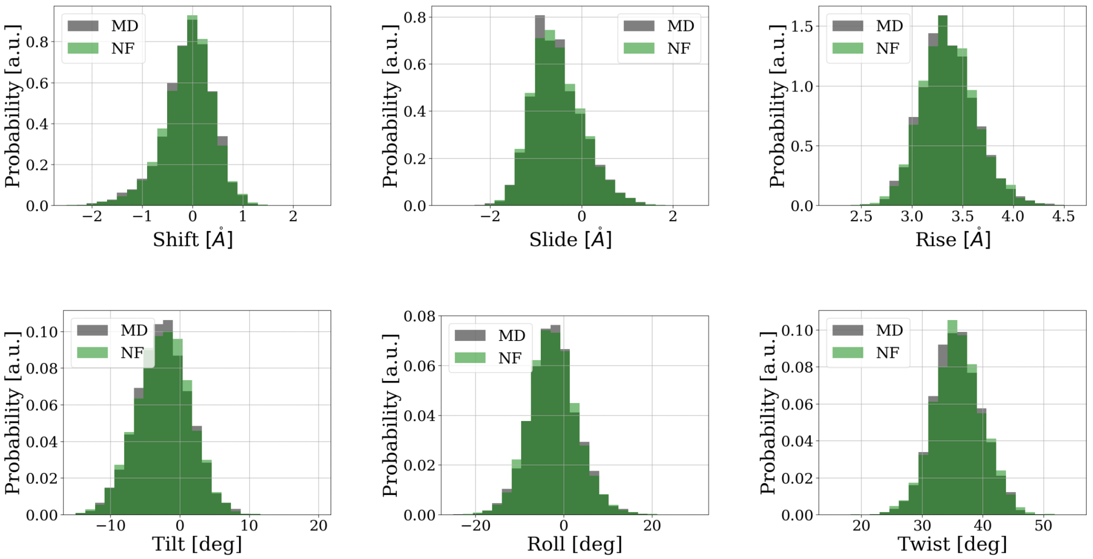


Figure 5: Distributions of the internal coordinates for the 5th base-pair step as sampled from atomistic MD simulations (grey) and the Normalizing Flow double-step model (green).

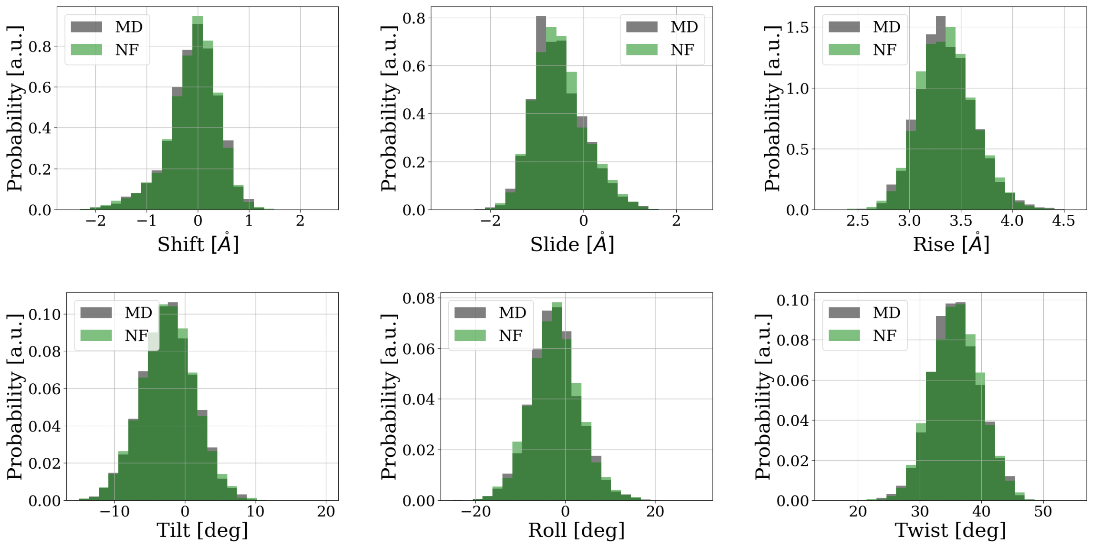

Figure 6: Distributions of the internal coordinates for the 5th base-pair step as sampled from atomistic MD simulations (grey) and the Normalizing Flow single-step model (green).

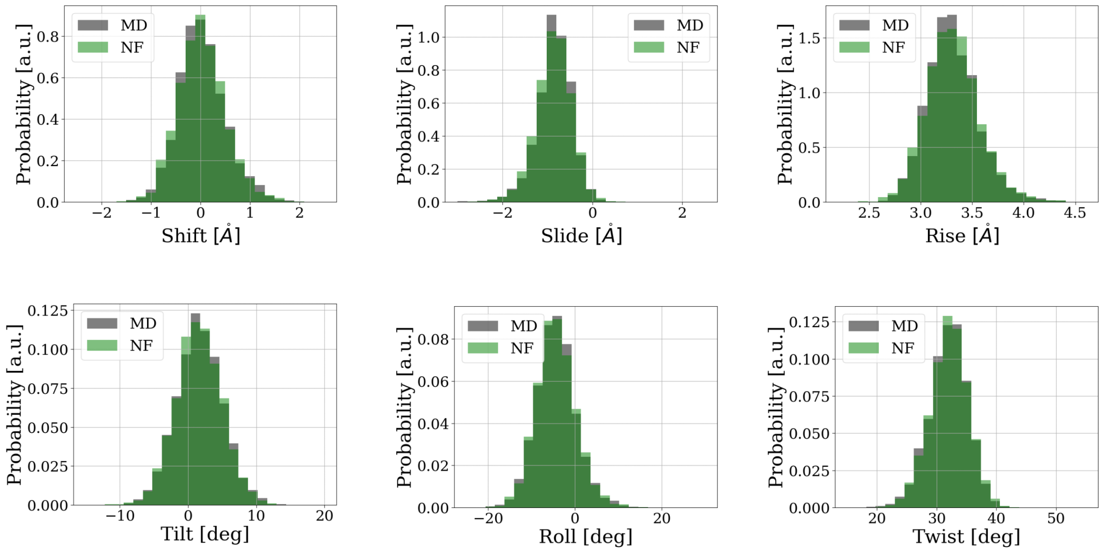

Figure 7: Distributions of the internal coordinates for the 6th base-pair step as sampled from atomistic MD simulations (grey) and the Normalizing Flow double-step model (green).

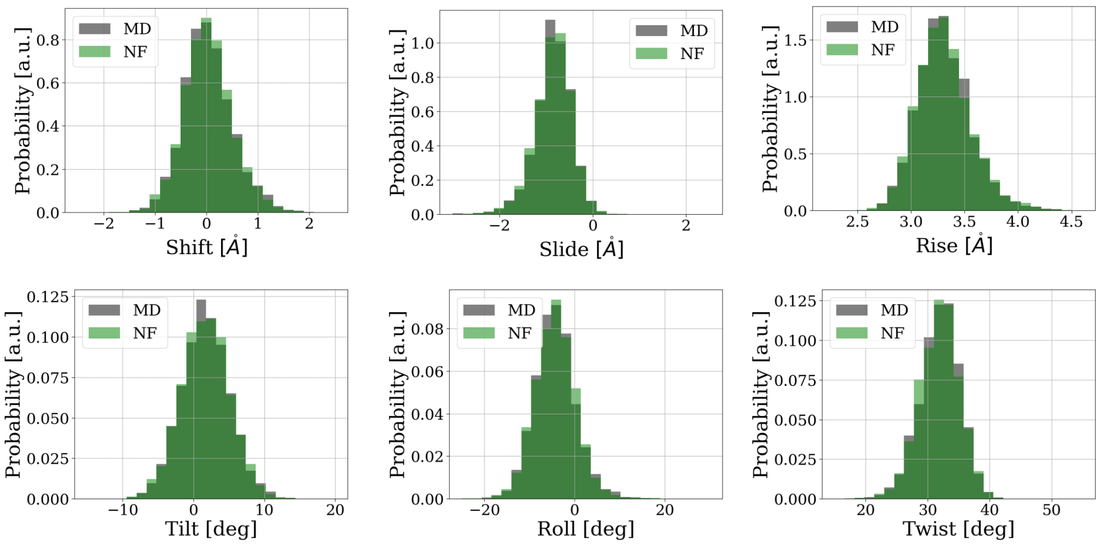


Figure 8: Distributions of the internal coordinates for the 6th base-pair step as sampled from atomistic MD simulations (grey) and the Normalizing Flow single-step model (green).

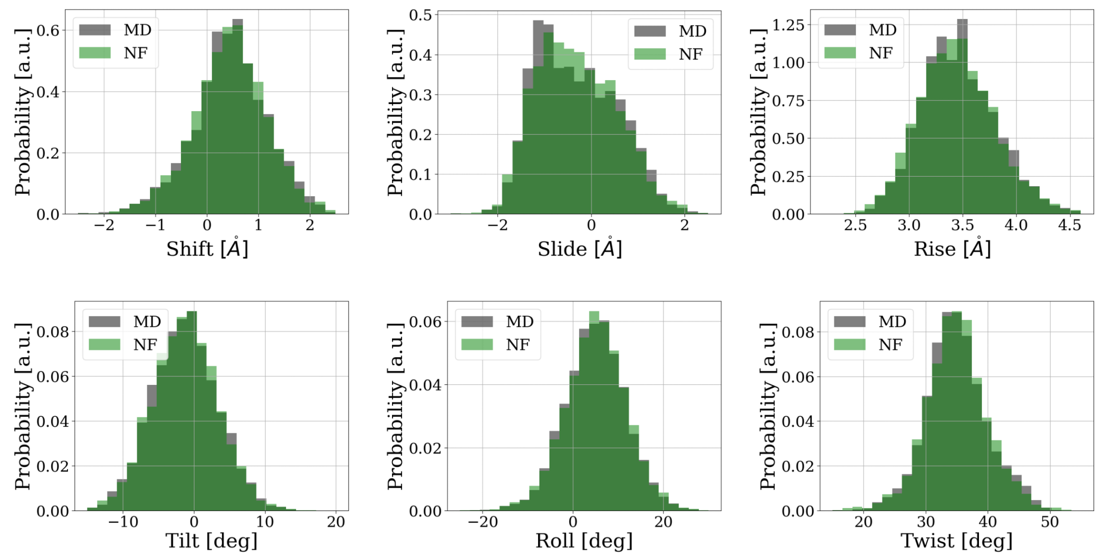


Figure 9: Distributions of the internal coordinates for the 7th base-pair step as sampled from atomistic MD simulations (grey) and the Normalizing Flow double-step model (green).

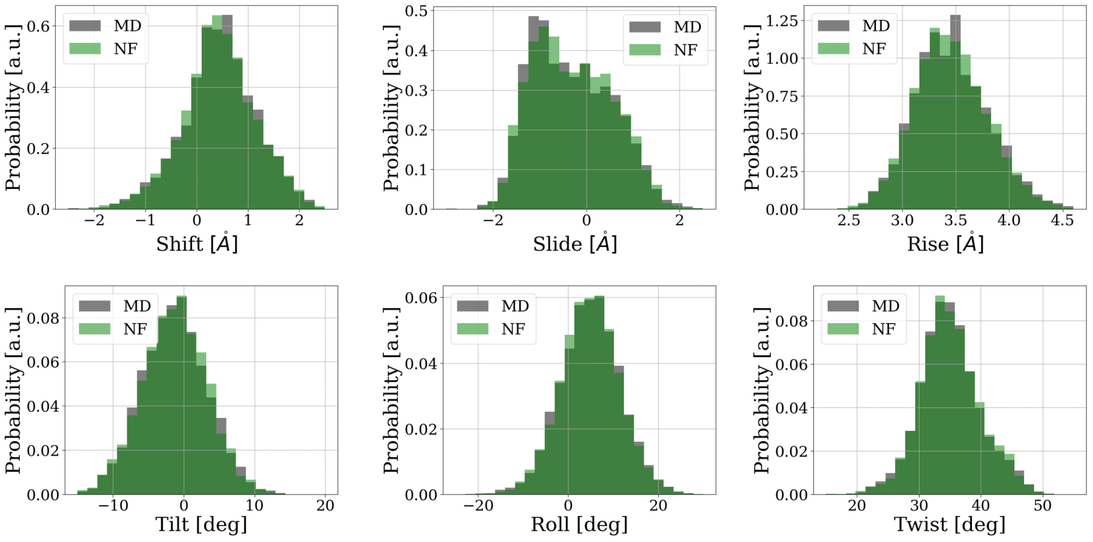


Figure 10: Distributions of the internal coordinates for the 7th base-pair step as sampled from atomistic MD simulations (grey) and the Normalizing Flow single-step model (green).

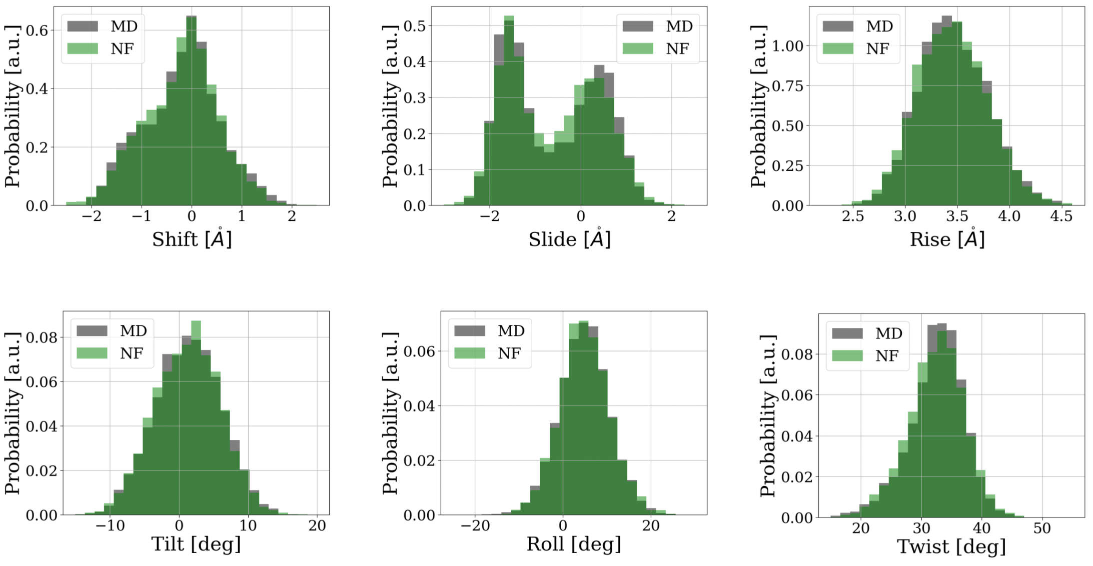


Figure 11: Distributions of the internal coordinates for the 8th base-pair step as sampled from atomistic MD simulations (grey) and the Normalizing Flow double-step model (green).

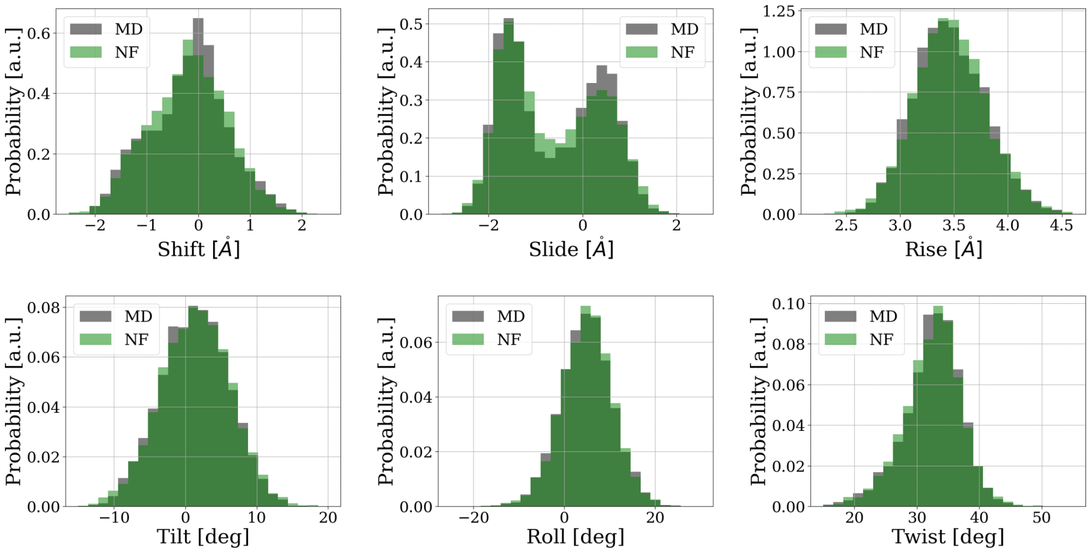


Figure 12: Distributions of the internal coordinates for the 8th base-pair step as sampled from atomistic MD simulations (grey) and the Normalizing Flow single-step model (green).

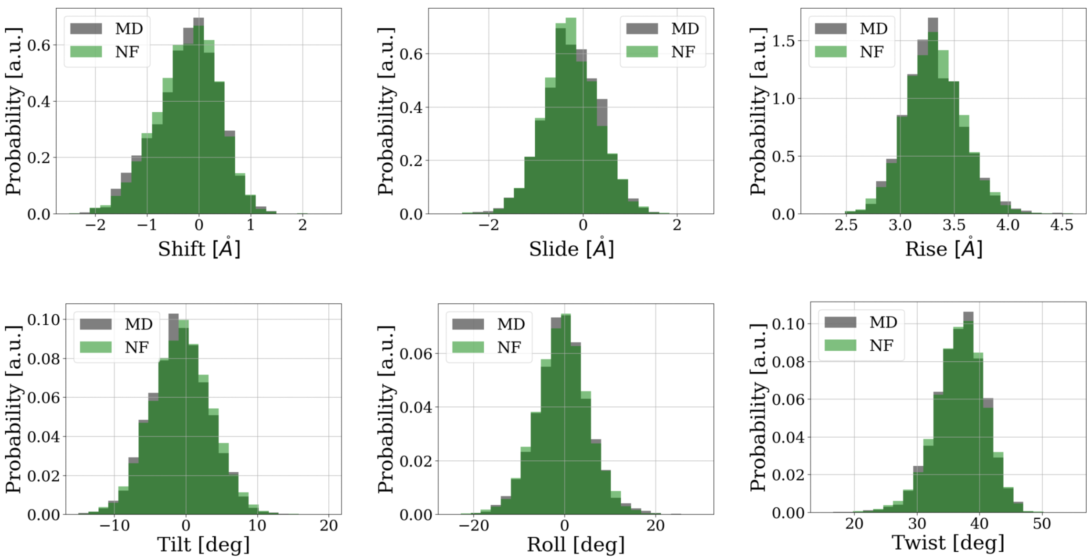


Figure 13: Distributions of the internal coordinates for the 9th base-pair step as sampled from atomistic MD simulations (grey) and the Normalizing Flow double-step model (green).

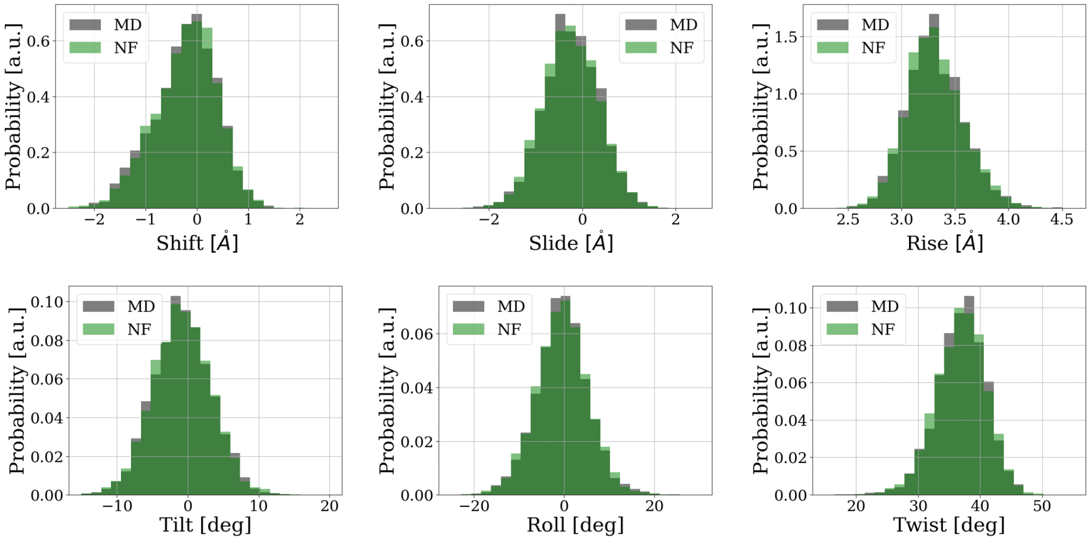


Figure 14: Distributions of the internal coordinates for the 9th base-pair step as sampled from atomistic MD simulations (grey) and the Normalizing Flow single-step model (green).

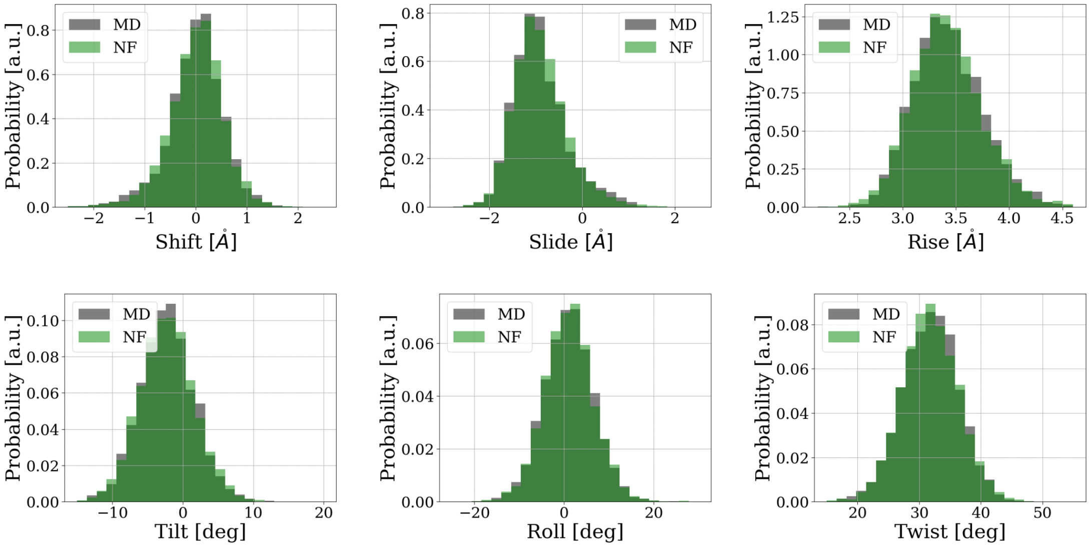


Figure 15: Distributions of the internal coordinates for the 10th base-pair step as sampled from atomistic MD simulations (grey) and the Normalizing Flow double-step model (green).

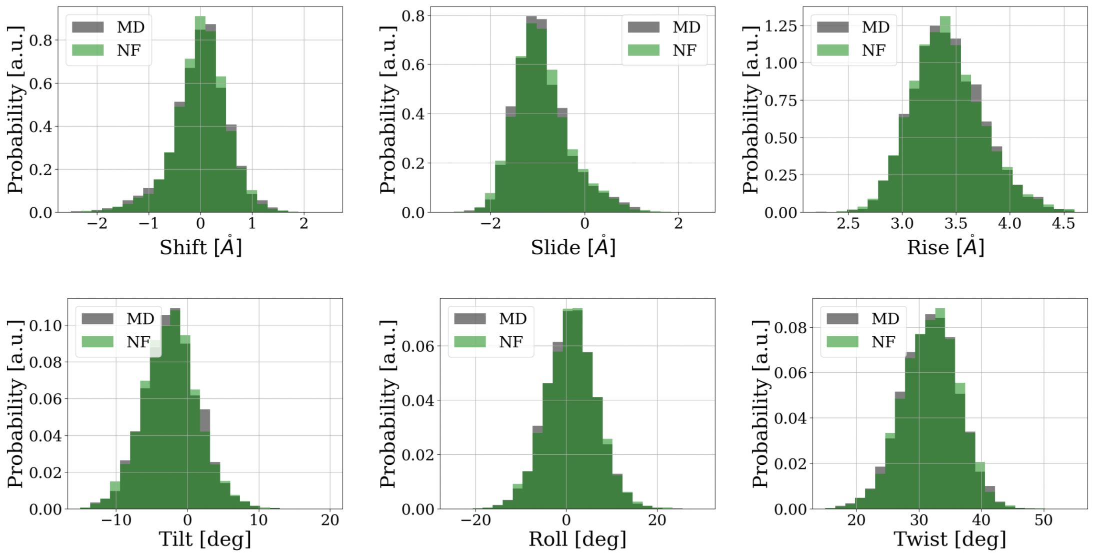


Figure 16: Distributions of the internal coordinates for the 10th base-pair step as sampled from atomistic MD simulations (grey) and the Normalizing Flow single-step model (green).

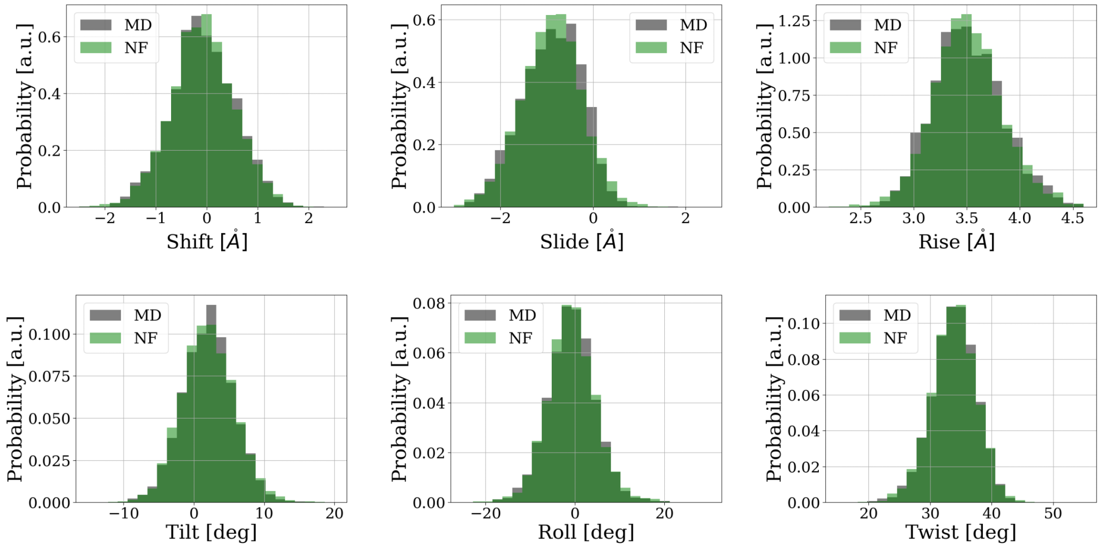


Figure 17: Distributions of the internal coordinates for the 11th base-pair step as sampled from atomistic MD simulations (grey) and the Normalizing Flow double-step model (green).

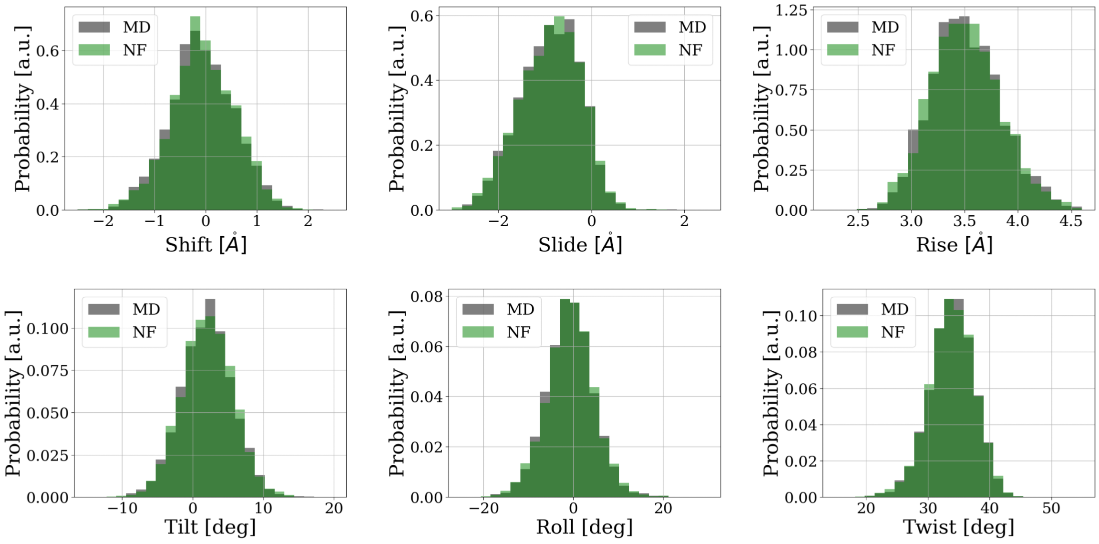


Figure 18: Distributions of the internal coordinates for the 11th base-pair step as sampled from atomistic MD simulations (grey) and the Normalizing Flow single-step model (green).

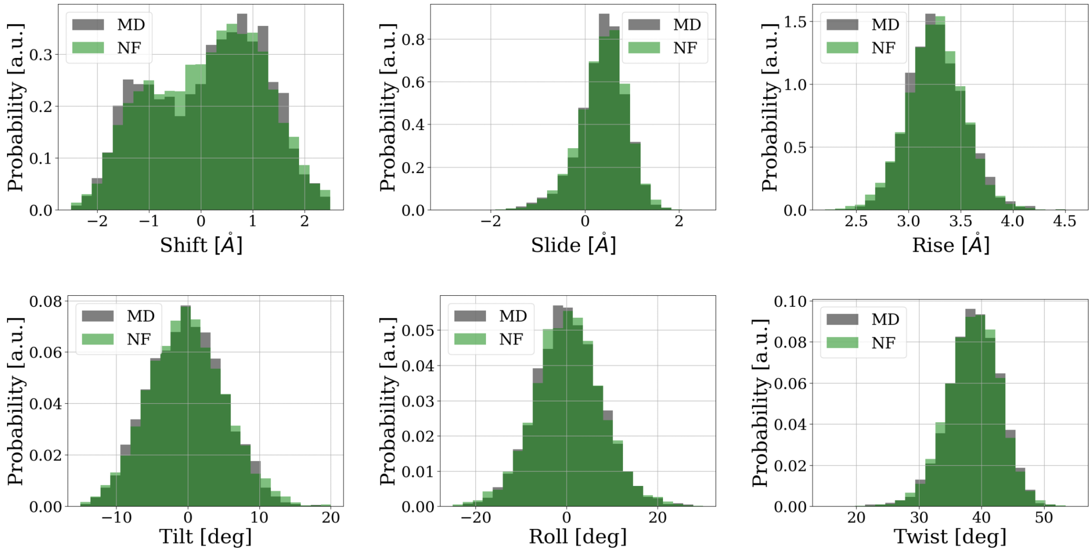


Figure 19: Distributions of the internal coordinates for the 12th base-pair step as sampled from atomistic MD simulations (grey) and the Normalizing Flow double-step model (green).

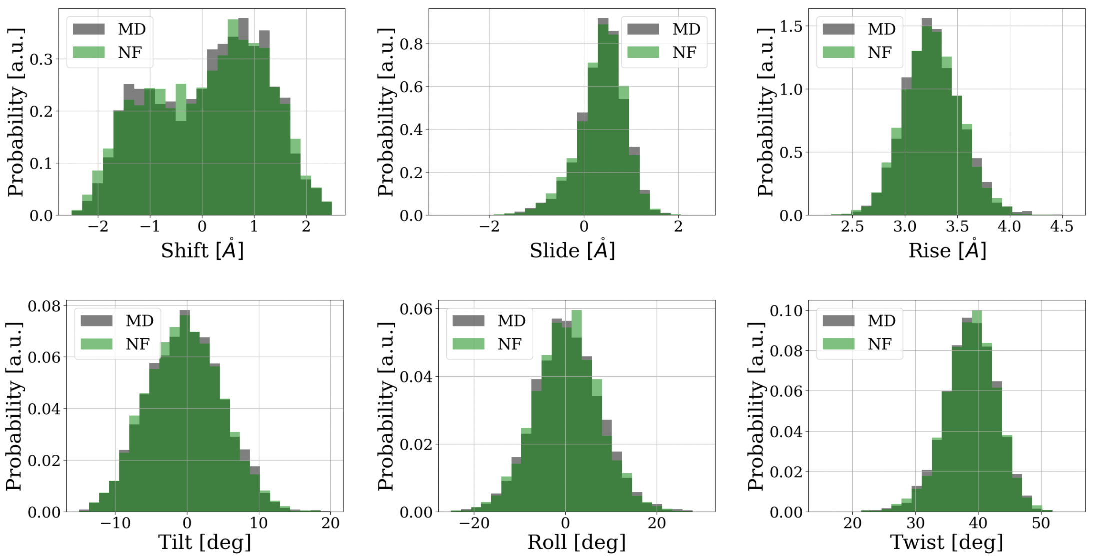


Figure 20: Distributions of the internal coordinates for the 12th base-pair step as sampled from atomistic MD simulations (grey) and the Normalizing Flow single-step model (green).

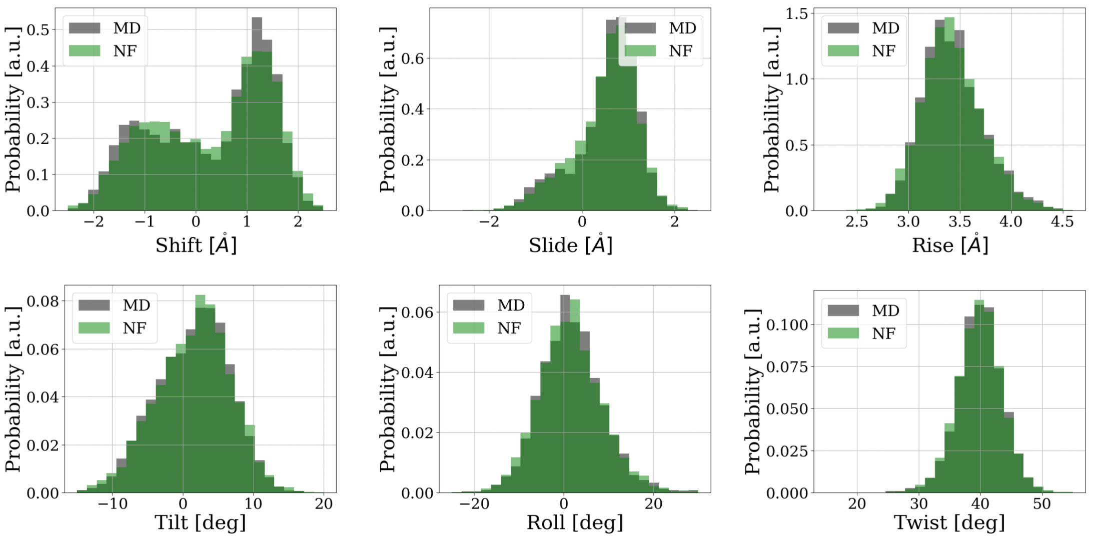


Figure 21: Distributions of the internal coordinates for the CA base-pair step within a CAA double-step as sampled from atomistic MD simulations (grey) and the Normalizing Flow double-step model (green).

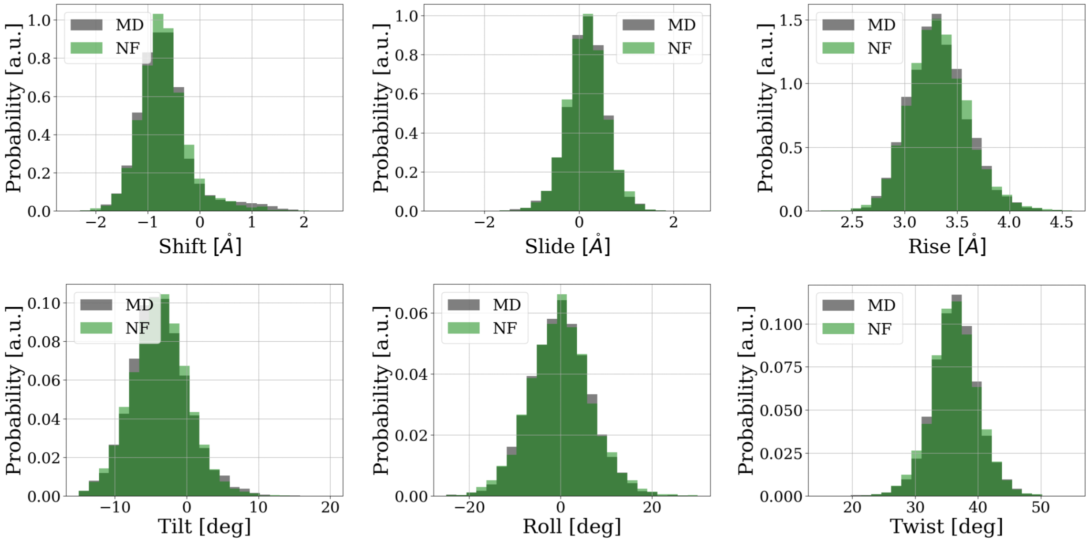


Figure 22: Distributions of the internal coordinates for the GA base-pair step within a GAG double-step as sampled from atomistic MD simulations (grey) and the Normalizing Flow single-step model (green).

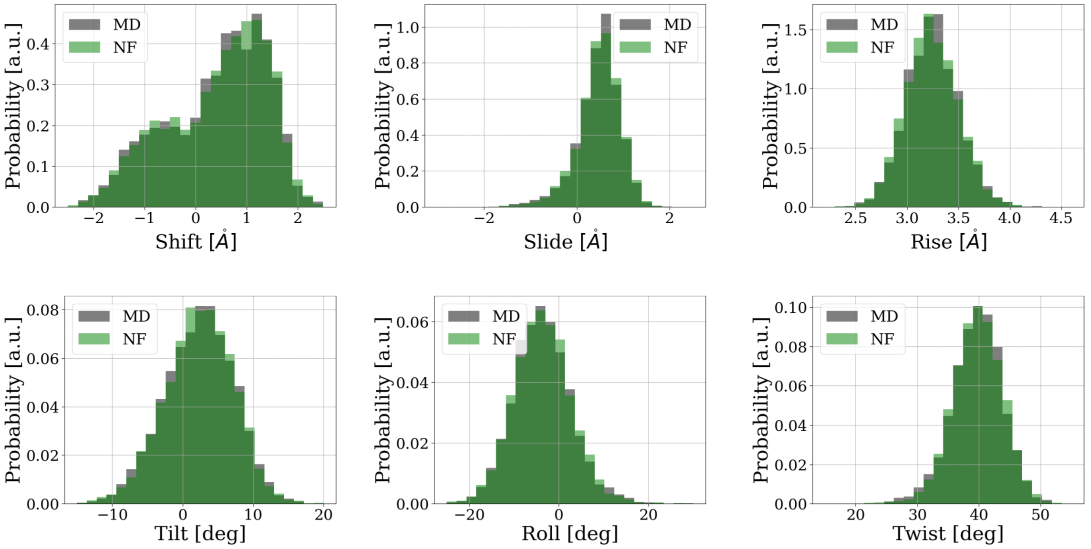


Figure 23: Distributions of the internal coordinates for the AC base-pair step as sampled from atomistic MD simulations (grey) and the Normalizing Flow single-step model (green).

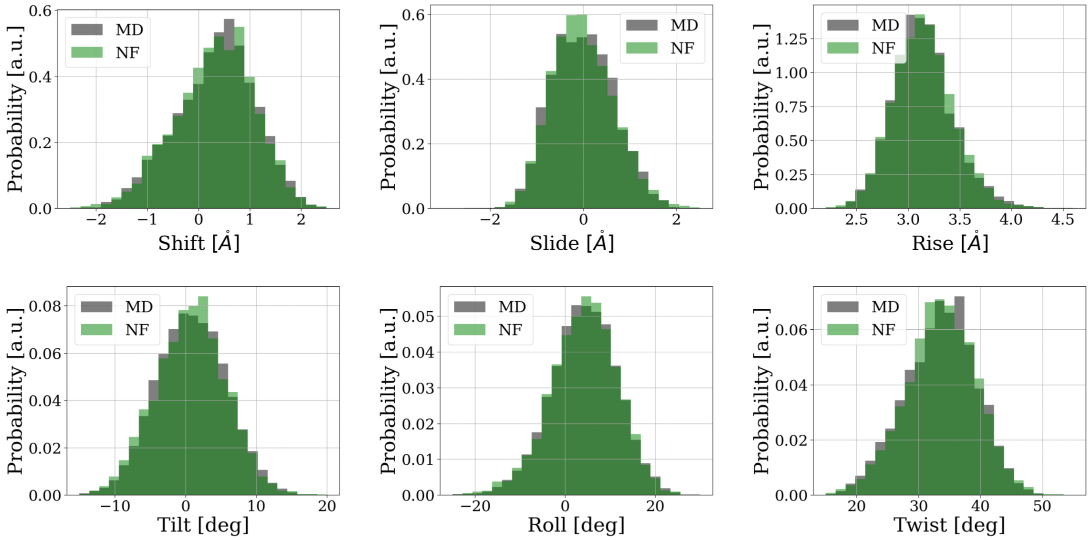


Figure 24: Distributions of the internal coordinates for the GG base-pair step as sampled from atomistic MD simulations (grey) and the Normalizing Flow single-step model (green).
